# Maggot : An ecosystem for sharing metadata within the web of FAIR Data

**DOI:** 10.1101/2024.05.24.595703

**Authors:** Daniel Jacob, François Ehrenmann, Romain David, Joseph Tran, Cathleen Mirande-Ney, Philippe Chaumeil

**Affiliations:** INRAE, Université de Bordeaux, UMR BFP, 71 av E Bourlaux, F-33140 Villenave d’Ornon, France; INRAE, Université de Bordeaux, UMR BIOGECO, 69 route d’Arcachon, F-33610 Cestas, France; European Research Infrastructure on Highly Pathogenic Agents (ERINHA AISBL), 98 rue du Trône B-1050 Bruxelles, Belgium; INRAE, Université de Bordeaux, Bordeaux Sciences Agro, ISVV, UMR EGFV, F-33140 Villenave d’Ornon, France; Université de Bordeaux, INRAE, UMR BFP, 71 av E Bourlaux, F-33140 Villenave d’Ornon, France

**Keywords:** FAIR, Data management, Metadata, Interoperability, Crosswalk, Controlled vocabulary, Ontologies, Thesaurus, Semantic artefacts

## Abstract

**Background:** Descriptive metadata are crucial for the discovery, reporting and mobilisation of research datasets. Addressing all metadata issues within the Data Management Plan often poses challenges for data producers. Organising and documenting data within data storage entails creating various descriptive metadata. Subsequently, data sharing involves ensuring metadata interoperability in alignment with FAIR principles. Given the tangible nature of these challenges, a real need for management tools has to be addressed to assist data managers to the fullest extent. Moreover, these tools have to meet data producers requirements and be user-friendly as well with minimal training as prerequisites.

**Results:** We developed Maggot which stands for Metadata Aggregation on Data Storage, specifically designed to annotate datasets by generating metadata files to be linked into storage spaces. Maggot enables users to seamlessly generate and attach comprehensible metadata to datasets within a collaborative environment. This approach seamlessly integrates into a data management plan, effectively tackling challenges related to data organisation, documentation, storage, and frictionless FAIR metadata sharing within the collaborative group and beyond. Furthermore, for enabling metadata crosswalk, metadata generated with Maggot can be converted for a specific data repository or configured to be exported into a suitable format for data harvesting by third-party applications.

**Conclusion:** The primary feature of Maggot is to ease metadata capture based on a carefully selected schema and standards. Then, it greatly eases access to data through metadata as requested nowadays in projects funded by public institutions and entities such as Europe Commission. Thus, Maggot can be used on one hand to promote good local versus global data management with open data sharing in mind while respecting FAIR principles, and on the other hand to prepare the future EOSC FAIR Web of Data within the framework of the European Open Science Cloud.

## Background

In the realm of scientific research, metadata plays a key yet often overlooked role. Despite their crucial importance for the discovery, reporting, and mobilisation of research datasets, metadata remains insufficiently known within scientific communities. Yet being data themselves, metadata have to be managed with the same level of rigour as the data produced and consumed by research processes. This lack of awareness persists at a time when the sharing of research data has emerged as a cornerstone of open or at least reproducible science initiatives. As the scientific landscape increasingly emphasises transparency and collaboration, understanding the significance of metadata becomes imperative [1, 2].

Producing comprehensive metadata is not a task to be taken lightly. It requires time, effort and expertise. Data producers tasked with generating datasets may be reluctant if they see no tangible return on investment in creating metadata [3]. To overcome this hurdle, proactive efforts are required to raise awareness among data producers about the benefits of open data practices [4]. However, crafting metadata poses challenges beyond incentivization. Data Management Plans (DMPs), which outline strategies for managing research data throughout their lifecycle, often pose non-trivial questions for data producers. These questions may be time-consuming or complex, particularly when datasets span diverse scientific domains and require input from individuals with varied skill sets. Consequently, collaborative efforts involving domain experts, data managers, and information specialists are essential for navigating the intricacies of DMPs effectively, furthermore when projects are involving multiple partners (e.g. [5]). The sheer diversity of research data and possible dimensions - i.e., the type of characteristics they describe - further complicate metadata management [6]. From omics data to images to experimental data tables, the spectrum of data types is vast and multifaceted.

Given the complexity of the matter, it is suitable to differentiate between various types and functions of metadata. While not delving into every category, we can simplify them into two main groups: general metadata and specialised metadata. Within the latter category, we encounter structural metadata, which serve to depict the organisation, arrangement, and interconnections within a dataset. For instance, when considering different types of data such as experimental data tables, it becomes obvious that structural metadata are essential for optimising their utility [7]. Conversely, general metadata (descriptive, administrative, rights) apply to all data types generated within studies with similar experimental contexts or even an entire project. The subsequent sections of this article will focus on these general metadata.

How can we collect such metadata while ensuring that they ultimately meet the requisite criteria for interoperability? Indeed, standardisation is the key towards interoperability and consistency in metadata practices. Sustainable metadata have to adhere to established standards and be described using controlled vocabularies endorsed broadly by the scientific community [8]. However, the responsibility for metadata creation predominantly falls upon data producers who possess intimate knowledge of the data intricacies. This is challenging as data producers may lack familiarity with metadata standards and best practices, and so reinforces the importance of roles within the data management ecosystem. Data managers and data stewards, equipped with expertise in applying FAIR (Findable, Accessible, Interoperable, Reusable) principles [9] and metadata standards, play a key role in guiding metadata creation and management processes. Conversely, scientists, serving as data producers, possess deep domain knowledge essential for contextualising and enriching metadata. Recognizing the complementary of these roles, collaborative partnerships between data managers and scientists are indispensable for ensuring the effective and sustainable management of research data [10].

Attempting to find a one-size-fits-all data warehouse capable of accommodating every data type proves to be a futile endeavour. Invariably, some data types remain unsupported or inadequately represented within existing data repositories. To address this challenge, we propose a pragmatic approach that involves managing data directly from storage spaces and then depositing them when the time comes into repositories tailored to requirements of each data type. To this end, we have developed Maggot (Metadata Aggregation on Data Storage), a specialised tool for aggregating metadata into data storage. It is specifically designed for annotating datasets by generating metadata files to attach to the data storage. It has been designed to help data managers in solving as many of the aforementioned challenges as feasible, all while catering to the needs of data producers with minimal training on their part.

### Design Considerations

The development and implementation of Maggot followed a structured approach, involving multiple steps and actors (**Fig. 1**). These steps encompass in particular the identification of metadata fields, terms, vocabularies, and standards. Inspiration for this approach was drawn from an online document detailing the implementation of a descriptive metadata plan [11]. An approach with some similarities has been adopted with the FAIR-DS application [12], dedicated to nucleotide sequences.

**Figure 1:**
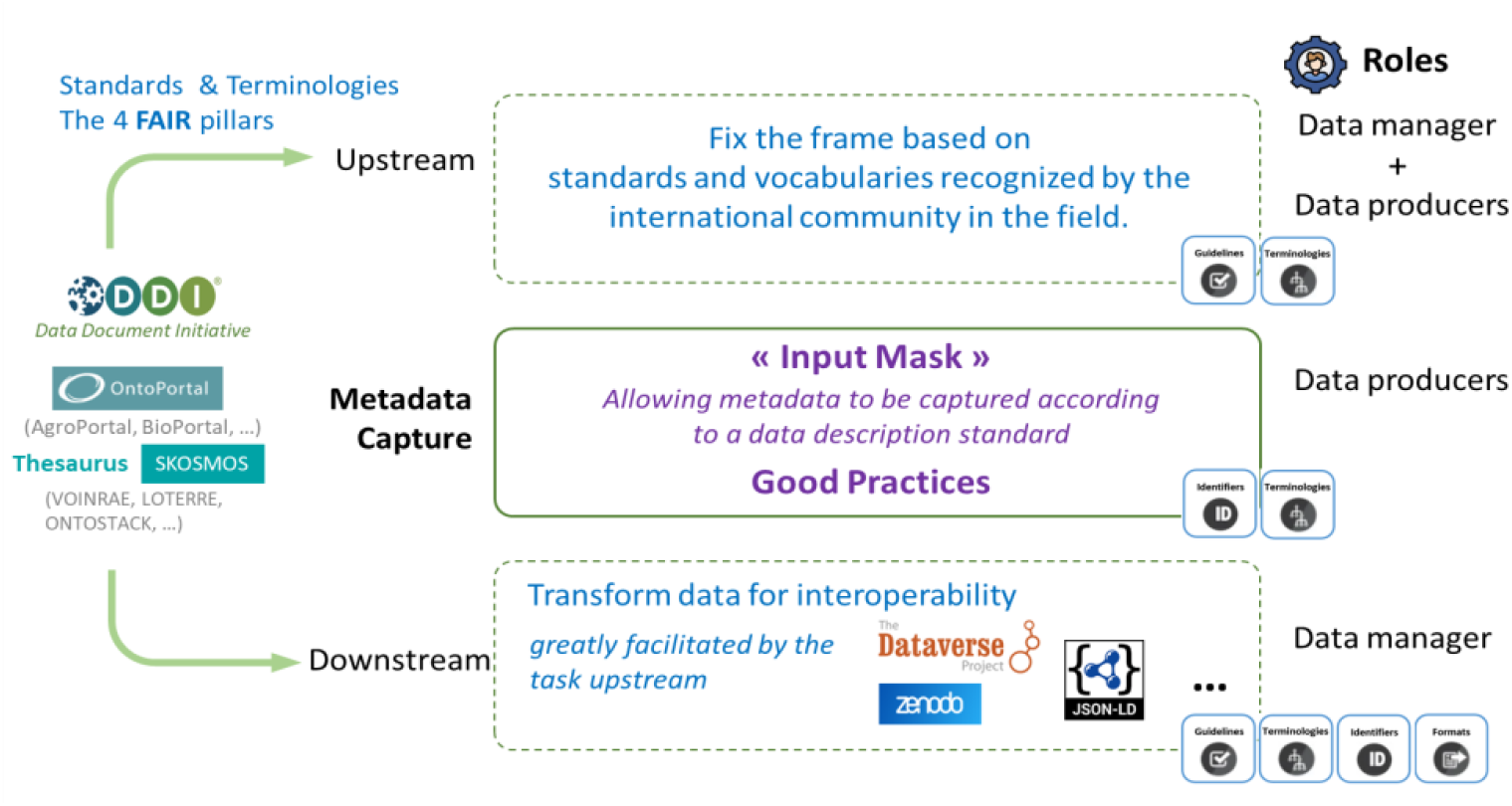
Development and implementation of Maggot tool structured into three steps, namely before, during and after metadata capture: i) Upstream, collaboration between the data manager and the data producers was essential to select and customise a flexible metadata schema adapted to the scientific domain as well as the identification of terms and vocabularies (dictionaries, thesauri, ontologies). Therefore, Maggot proposes the Dataverse schema serving as a fundamental model, itself based on the standard DDI metadata schema. ii) For metadata entry, data producers must be trained on good practices such as the proper use of permanent identifiers or the choice of licences. iii) Downstream, data can easily be pushed into a support data repository without any addition or can be harvested based on a dedicated protocol (OAI-PMH). JSON-LD format is also supported for linking metadata within the realm of linked data, thereby ensuring their interoperability. The complementarity of roles between data manager and data producers ensures effective and sustainable research data management.

Within any research team, collaboration between the data manager and data producers is essential to select and customise the minimum metadata required and an associated metadata schema suitable for the specific scientific domain. While this process may present challenges, it is crucial to meticulously construct and adapt the schema to align with existing data. Rather than mandating data conformity, the schema should be flexible to accommodate pre-existing data. Thus, the adoption of a schema, such as the one implemented within the Harvard Dataverse software [13] (https://dataverse.harvard.edu), followed by an iterative and progressive adjustment, is the approach embraced by Maggot. Indeed, Harvard Dataverse itself is built upon the standard DDI (Document, Discover and Interoperate) metadata schema (https://ddialliance.org), which has been expanded to accommodate its requirements. The advantage of the DDI schema is that it encompasses a wealth of general information for describing datasets of any type.

In the same way, Maggot advocates an iterative and progressive approach regarding the management of controlled vocabulary. Recognizing the impossibility of achieving exhaustiveness in the initial stages, Maggot facilitates a process of continuous improvement. This involves starting with a simple vocabulary dictionary sourced locally and consolidating community-used vocabulary within the related scientific domain. Subsequently, consideration could be given to the creation of a thesaurus (or at least a controlled vocabulary), with or without mapping to existing ontologies. Maggot is seamlessly based on the SKOSMOS web application [14] (https://skosmos.org) to query thesauri directly, streamlining the process. Furthermore, ontologies can be chosen progressively by selecting those which are truly relevant for the collective and by drawing up an understandable landscape of the context in which they fit. In the same way for thesauri, Maggot offers the opportunity to enrich metadata using ontologies seamlessly accessible through the OntoPortal web applications such as BioPortal [15] (https://bioportal.bioontology.org) or AgroPortal [16] (https://agroportal.lirmm.fr).

In our view, creating metadata should not compel data producers to engage in training in topics or concepts outside their expertise, such as FAIR principles, the semantic web, or metadata schema, unless they choose to do so. The data manager needs to recognize that data producers may not possess extensive knowledge of data management practices, especially in the realm of open science and FAIR data. Therefore, pedagogy should be prioritised by refraining from overwhelming data producers with technical details specific to their field. Instead, the data manager should focus on raising awareness and encouraging data producers to improve the quality and reusability of their data [3]. This includes providing guidance on relevant metadata and controlled vocabulary within their scientific domain, as well as training on best practices such as the use of permanent identifiers like DOI, ORCID, RoR, ePICs, Handle and other well recognized identifying systems. Furthermore, data producers should be informed about the selection of appropriate licences, such as CC-BY licences (https://creativecommons.org). One of the features of Maggot is precisely to allow the data manager to document each of the terms, providing examples and links for additional information if necessary. This contextual online assistance is thus accessible during entry to allow data producers to fill in each field in the most relevant way, ensuring optimal support throughout the process.

For downstream metadata management, Maggot provides functionality enabling transformation of metadata in two ways: to targeted data repositories with prerequisites as defined by the Core Trust Seal [17] (https://www.coretrustseal.org), or to an export format suitable for harvesting data by third-party applications via an application programming interface (API). These functionalities are built upon a metadata crossing approach based on the mapping defined upstream by the data manager. They ensure compatibility with systems allowing content indexing, thus aligning with the FAIR principles. Adopting this approach improves organisations’ data-management practices by effectively using metadata throughout the data lifecycle and facilitates data linkage. It enhances data interoperability and reusability, optimising the value derived from data assets. It also provides practical accessibility through the Web of FAIR Data - i.e. data which meet FAIR principles - by increasing data linkage possibilities, as envisioned by international consortiums like the European Open Science Cloud (EOSC, https://open-science-cloud.ec.europa.eu).

Finally, let us emphasise that a software application rarely meets a need alone but is part of a more global approach, involving several roles. Namely, the data manager is the person who defines the data policy, i.e., its implementation and governance, while data stewards are responsible for data quality, and therefore have a role in data curation. Unavoidably, the development and dissemination of metadata, involving a metadata-centric culture, underline the need for ongoing training initiatives and data stewards are pivotal in educating data producers. Despite advancements in tool intuitiveness and automation, the growing complexity and scale of data across the Web of Data call for an increase in the number of skilled data stewards within research organisations to ensure maximum data reusability [18].

## Results

Maggot (Metadata Aggregation on Data Storage) was developed to address the need for a multi-purpose data management tool capable of supporting the wide data diversity range within a collective. Its primary objectives are to provide visibility into the collective’s data assets, facilitate better data description and increase early adoption of FAIR principles. It contributes to guarantee the sustainability of data reusability, especially for those produced by fixed-term personnel (limited-term contract, doctoral students and postdoctoral fellows). It helps to raise awareness among newcomers and students about the importance of robust data description practices which is crucial for fostering a culture of data management excellence [3].

Maggot enables users to achieve these objectives through various features and functionalities which can be divided into three parts: creation, sharing, dissemination (**Fig. 2**). Maggot supports by default the common metadata standards of Harvard Dataverse (based on DDI) serving as a starting point which can be extended/specialised to suit individual contexts. It then offers scalability and flexibility to enrich the core metadata to adapt any experimental context. Controlled vocabulary management is another key aspect of Maggot, offering options ranging from simple dictionaries to ontologies as discussed above. The tool includes enrichment functionalities of existing resources (e.g., dictionary editing, adding additional ontologies), ensuring that users can effectively manage and utilise controlled vocabularies relevant to their data. To this end, it offers great flexibility in configuration, allowing organisations to tailor the tool to their specific needs and requirements (**Fig. 3**).

**Figure 2:**
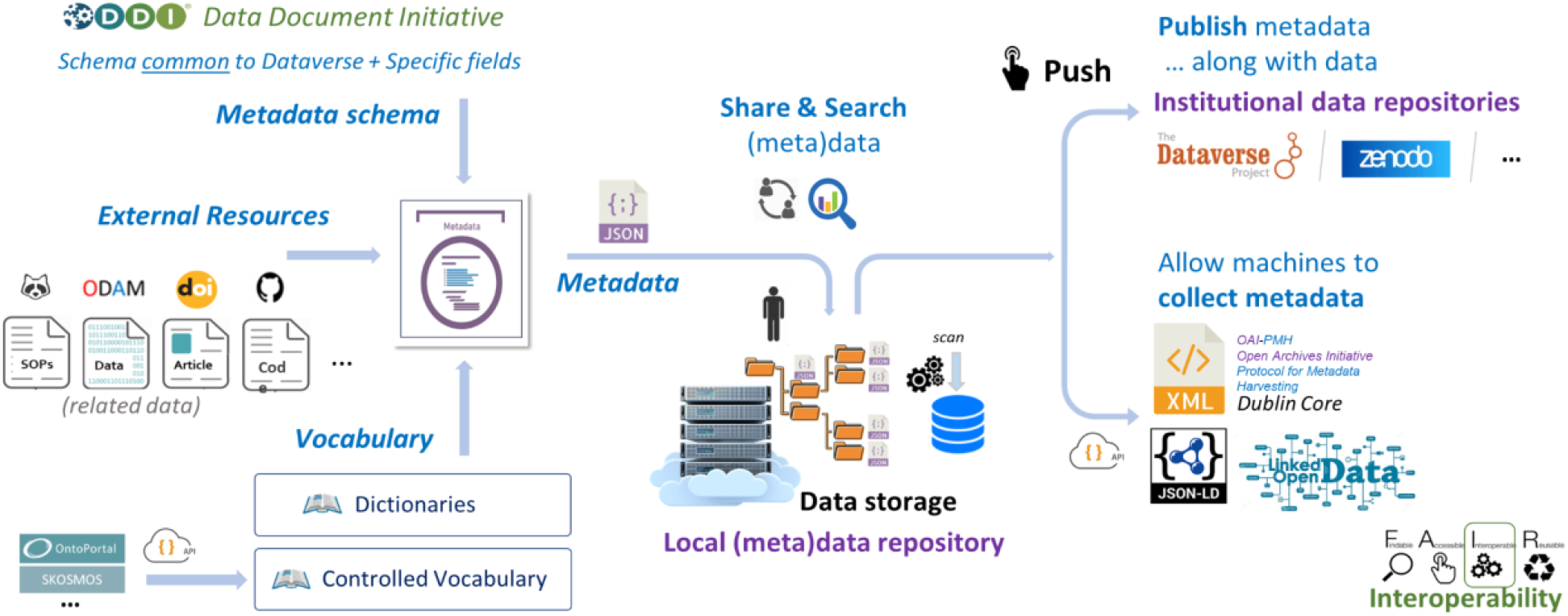
Main functionalities of Maggot split into three parts: creation, sharing, dissemination. First, producing a document with metadata sets of data within a collective of people, thus allowing i) to answer certain questions of the Data Management Plan (DMP) concerning data organisation, documentation, storage and sharing in the data storage space, ii) to meet certain data and metadata requirements, listed for example by Open Research Europe in accordance with FAIR principles. Next, searching for datasets by their metadata. Indeed, the descriptive metadata thus produced can be associated with the corresponding data directly in the storage space and then it is possible to perform a search on the metadata to find one or more sets of data. Only descriptive metadata is accessible by default. Finally, publishing the metadata of the datasets as well as their data files in a European-approved repository, with the possibility either to directly harvest the metadata via the OAI-PMH protocol, or to export the associated metadata with their semantic context for full interoperability.

**Figure 3:**
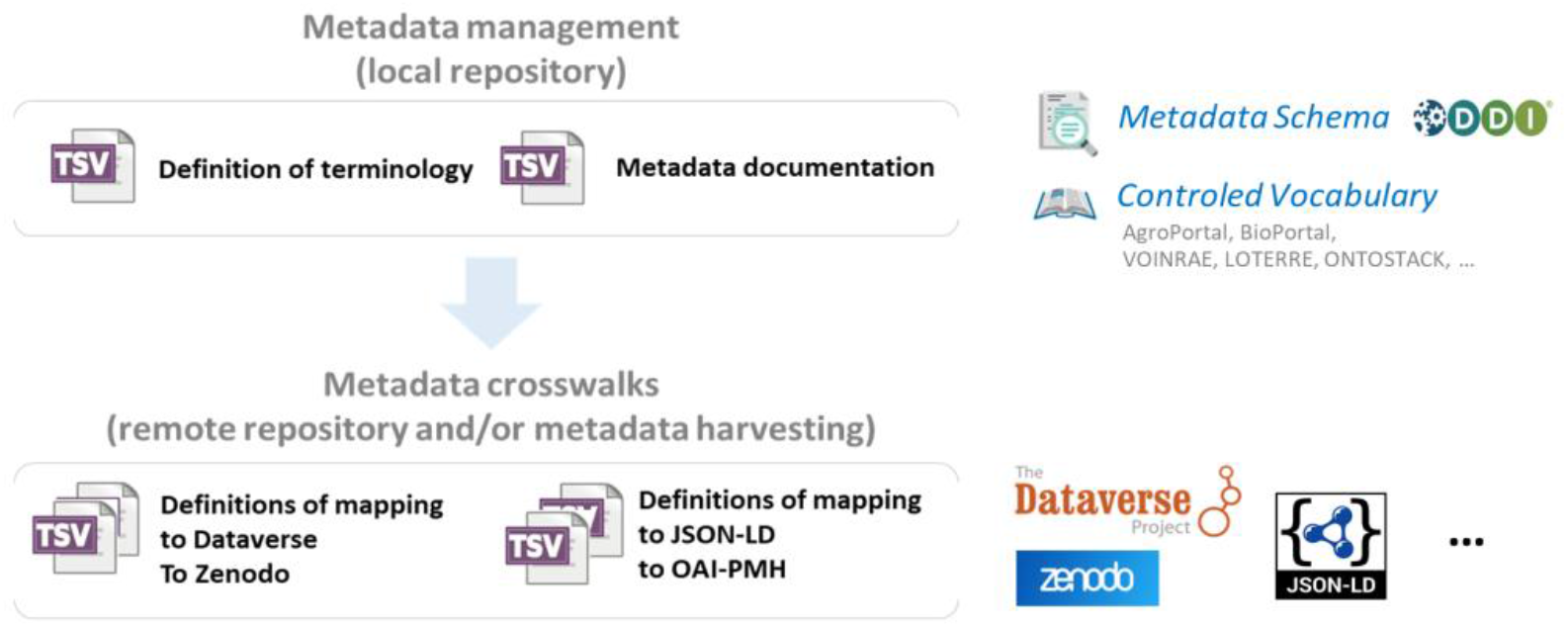
Maggot tool flexibility in configuration. Maggot allows users to choose all the metadata describing their data with two levels of definition files. The first level concerns the definition of metadata similar to a descriptive metadata plan. This category is more akin to configuration files, and constitutes the heart of the configuration around which everything else is based. The input and search interfaces are completely generated from these definition files, thus defining each of the fields, their input type and the associated controlled vocabulary. The second level concerns the definitions of the mapping to a differently structured metadata schema (metadata crosswalk, i.e a specification for mapping one metadata standard to another), used either for metadata export to a remote repository (e.g. Dataverse, Zenodo) or for metadata harvesting (e.g. JSON-LD, OAI-PMH).

Implementing a data management plan (DMP) entails certain prerequisites, including data externalisation to preserve them outside users’ disk space. This ensures data is secured in one location and serves as an initial backup, which becomes particularly crucial when fixed-term personnel are involved in data production. Consequently, considerations arise regarding the organisation of storage spaces, such as harmonising folder and file naming conventions, setting up folder structure, and using README files to provide essential information. Maggot precisely answers these DMP challenging questions related to organising, documenting, storing and sharing data from various sources. In our approach the data storage space becomes a local reference data repository, mitigating the risk of data duplication or divergence to another medium. Then, only metadata need to be added to this centralised space, streamlining data management processes, and enhancing efficiency. Indeed, once the metadata file is generated in JSON format, it has to be placed in the storage space reserved for this purpose alongside the corresponding dataset. This metadata file can be seen as a README file adapted to machines (**Additional file** 1), but still readable by humans. In contrast, with an internal structure, it offers better coherence and consistency of information than a simple README file with a completely free and therefore unstructured text format. In this way, the storage space becomes a data asset which can therefore be efficiently leveraged using metadata. Indeed, all the JSON metadata files are scanned and parsed according to a fixed time interval (30 min) then loaded into a database. This allows users to query datasets based on metadata filters. The search form, in a compact shape, is almost the same as the entry form. Matching datasets are returned as a list, and for each of them a provided link helps to access the detailed metadata.

Metadata entry can be initiated at the outset of a project or study, without requiring the completion of all data acquisition or processing, or waiting until the data need to be published. Indeed, the ability to reload a metadata file facilitates gradual and iterative metadata addition across the project, thereby spanning the research data lifecycle to the greatest extent possible. Maggot supports the input of both descriptive and administrative metadata for any type of data, including datasets, images, sequences, and more, with customizable field definitions to suit diverse user requirements. Moreover, Maggot emphasises ease of use and adaptability. It offers guided assistance through drop-down menus and vocabulary lists featuring autocompletion, greatly speeding up the process of filling in numerous descriptive metadata. Crucially, Maggot does not restrict the choice of data repository, ensuring compatibility with currently supported platforms knowing that others may be supported in the future (e.g. Dryad [19], RO-Crate [20]). This also does not prejudge the use of metadata. It is entirely possible, for example, to set up an internal metadata harvesting process to automatically fill in another data source (e.g., FAIRDOM-SEEK data management platform [21]). It is essential to highlight that opting for Maggot to generate metadata does not confine the data to an isolated silo. In case one day the Maggot tool was no longer supported, all metadata will persist in disk space in a format accessible to both humans and machines. This ensures that future applications/services are able to continue to use legacy metadata and therefore warranty data reuse. For this purpose, Maggot enables data scientists or data repositories to harvest data. The OAI-PMH (Open Archives Initiative Protocol for Metadata Harvesting, https://www.openarchives.org) allows for listing all datasets based on the DublinCore schema (https://www.dublincore.org), while the metadata of each dataset can be harvested in JSON-LD format (JSON for Linking Data, https://json-ld.org), mainly adhering to the schema.org standard (https://schema.org). This aspect is particularly critical for linking metadata within the realm of linked data, thereby ensuring their interoperability. For instance, we plan in the near future to support DCAT-based harvesting (https://www.w3.org/TR/vocab-dcat-3/).

Maggot also provides a solution to data fragmentation. Indeed, data is often scattered across various platforms, databases, and file formats, making it challenging to locate and access. Moreover, non-standardized metadata and inconsistent data organisation hinder effective data discovery and reuse. Therefore, Maggot allows data producers to specify resources, i.e., data in the broader sense, whether external or internal, to centralise all links towards data (**Fig. 2**). External resources must be specified by an URL with preference for a permanent identifier (e.g., DOI) but also any URL pointing to data whether they comply with the FAIR principle or not. Furthermore, in the case of local data management, it is wise to indicate in which space the data is located if it is not located in the same place as metadata (e.g., NAS unit or data cloud). Maggot can thus become a data hub by gathering all references to several data sources in one place at hand.

While the primary focus is managing metadata linked to data stored within a given collective, Maggot also facilitates data openness through metadata, especially in projects funded by public institutions. By setting up metadata schemas that facilitate crosswalks with established data repositories, users can seamlessly push metadata along with the corresponding data without the need for additional data entry. This promotes data openness and accessibility in accordance with international standards and community norms. Such functionality empowers organisations to share their data with external stakeholders while ensuring consistency and interoperability. Maggot thus offers a comprehensive and open solution for metadata management, catering to the diverse requirements of organisations and promoting best practices in data description and dissemination.

### Implementation and Documentation

The deployment of Maggot requires two infrastructure components: a dedicated server for the web application and a designated data storage space. Regarding the server, it must be capable of running an operating system compatible with Linux. In addition, it should support containerization using Docker. This latter aspect offers a simplified approach to installation and administration, but also ease of use and flexibility. Regarding data storage, any technology is suitable. Data storage can be local (e.g., NAS unit) or remote (e.g., data cloud). Successful tests have been performed by implementing a server on our institutional data center and data storage on another data center. Access to the storage space can easily be done using the rclone tool (https://rclone.org), a real Swiss army knife for disk sharing.

Maggot is a web-based PHP application as a front-end using MongoDB (https://www.mongodb.com) to index all scanned metadata from disk storage every 30 minutes. Moreover, Maggot mobilises various vocabularies (thesauri and ontologies), most of which are in remote resources. So the utilisation of APIs plays a significant role, particularly for integrating these vocabularies. This extensive use of APIs facilitates real-time imports, thus reducing the need for pre-updating information.

Documentation is available via https://inrae.github.io/pgd-mmdt/ and from within the app. It includes technical information on how to configure Maggot, but also a quick overview of how to use it. For data managers, it explains in detail how to construct the terminology with the associated vocabularies.

## Conclusion

Maggot is specifically designed to annotate datasets by generating metadata files to be linked into storage spaces, to tackle challenges related to data organisation, documentation, storage, and frictionless FAIR metadata sharing within the collaborative group and beyond. Indeed, Maggot meets the Open Data requirements beyond the simple provision of data with unlimited access. This essentially implies: i) to ensure search and access to metadata that define data access and usage conditions, and ii) to foster metadata and data interoperability to break down silos, highlighting the necessity of embracing FAIR principles even when complete openness is not achievable.

By covering as much of the research data lifecycle as possible, Maggot ensures effective and sustainable research data management and significantly simplifies the adoption of FAIR principles thereby empowering organisations to elevate the value and usability of their own data assets. Moreover, its ability via crosswalk approaches to distribute metadata based on standard schemas while being machine-readable, expands the toolbox needed to prepare the future EOSC FAIR Web of Data within the framework of the European Open Science Cloud.

### Availability of Source Code and Requirements

- Project name: Maggot
- Project homepage: https://pmb-bordeaux.fr/maggot/
- Project code repository: https://github.com/inrae/pgd-mmdt
- Documentation: https://inrae.github.io/pgd-mmdt/
- Operating system(s): Platform independent
- Programming languages: PHP, python, javascript
- Licence: GNU GPL v3
- Maggot, RRID: SCR_025261
- Biotools: https://bio.tools/maggot

## Supporting information

Examples of metadata files along with corresponding definitions files within an Excel workbook

## Abbreviations

API: Application Programming Interface
DDI: Document, Discover and Interoperate
DMP: Data Management Plan
EOSC: European Open Science Cloud
FAIR: Findable, Accessible, Interoperable, Reusable
JSON: JavaScript Object Notation
JSON-LD: JSON for Linking Data
OAI-PMH: Open Archives Initiative Protocol for Metadata Harvesting
NAS: Network Attached Storage

## Authors’ Contributions

Conceptualization: D.J, F.E, PC.; funding acquisition: D.J, F.E.; methodology: D.J., F.E, R.D.; software: D.J, F.E.; writing—original draft: D.J., R.D.; writing—review and editing: All authors. All authors read and approved the final manuscript.

## Acknowledgments

We thank Dr Catherine Deborde (INRAE UR BIA & PROBE Research Infrastructure, BIBS Facility Nantes) and Dr Annick Moing (INRAE UMR 1332 BFP, Bordeaux) for advice on the manuscript and for constructive reviews. We also thank Edouard Guitton (INRAE, Animal Health Department) for his fruitful feedback in the implementation of Maggot within his research infrastructure.

## Additional files

**Additional file 1** : Examples of metadata files along with corresponding definitions files within an [Excel workbook](https://docs.google.com/spreadsheets/d/1jGJDMKVKF-W0eWq3NdPHJzmps-wBXt5t/edit#gid=1815643208).

